# Application of CNTF or FGF-2 increases the number of M2-like macrophages after optic nerve injury in adult *Rana pipiens*

**DOI:** 10.1101/494831

**Authors:** Rosa E. Blanco, Giam S. Vega-Meléndez, Valeria De La Rosa-Reyes, Clarissa del Cueto, Jonathan M. Blagburn

## Abstract

We have previously shown that a single application of the growth factors ciliary neurotrophic factor (CNTF) or fibroblast growth factor 2 (FGF-2) to the crushed optic nerve of the frog, *Rana pipiens*, increases the numbers and elongation rate of regenerating retinal ganglion cell axons. Here we investigate the effects of these factors on the numbers and types of macrophages that invade the regeneration zone. In control PBS-treated nerves, many macrophage-like cells are present 100 μm distal to the crush site at 1 week after injury; their numbers halve by 2 weeks. A single application of CNTF at the time of injury triples the numbers of macrophages at 1 week, with this increase compared to control being maintained at 2 weeks. Application of FGF-2 is equally effective at 1 week, but the macrophage numbers have fallen to control levels at 2 weeks. Immunostaining with a pan-macrophage marker, ED1, and a marker for M2-like macrophages, Arg-1, showed that the proportion of the putative M2 phenotype remained at approximately 80% with all treatments. Electron microscopy of the macrophage-like cells at 1 week shows strong phagocytic activity with all treatments, with many vacuoles containing axon fragments and membrane debris. At 2 weeks with PBS or FGF-2 treatment the remaining macrophage-like cells are less phagocytically active, containing mainly lipid inclusions. With CNTF treatment, at 2 weeks many of the more numerous macrophages are still phagocytosing axonal debris, although they also contain lipid inclusions. We conclude that the increase in macrophage influx seen after growth factor application is beneficial for the regenerating axons, probably due to more extensive removal of degenerating distal axons, but also perhaps to secretion of growth-promoting substances.

## Introduction

Central nervous system (CNS) neurons react to injury in different ways in different groups of animals – for example, mammalian neurons mostly die and the remainder show poor regrowth [1,2], fish CNS neurons survive and regenerate successfully [3,4], while amphibian neurons show intermediate survival rates and successful regrowth [5]. Neuronal death after injury is thought to be due to interruption of the supply of neurotrophic factors from the target region [1,6–8], while poor regrowth is due to an inhibitory environment [9,10], and many studies have been devoted to alleviating these conditions.

In recent years, it has been proposed that macrophages play a key role in modulating the progression of neurodegenerative diseases [11–14] and also the response to CNS injury [15–17]. Macrophages originate from bone-marrow-derived monocytes, which circulate in the bloodstream [18] and are then capable of infiltrating injured tissues, where they differentiate into macrophages [19,20]. Along with already-resident microglia, these cells phagocytose debris [21] and secrete chemicals that enhance or inhibit the inflammatory response [22,23]. However, the potential roles of phagocytic cells in modulating neuronal survival and axonal regrowth after injury remain somewhat ambiguous, in part because of their dual pro- and anti-inflammatory phenotypes [24,25].

Our previous work has concentrated on retinal ganglion cell (RGC) survival after damage to the optic nerve of the frog, *Rana pipiens*, and the beneficial effects of topical growth factor administration upon that survival [26–30]. Recently we showed that the speed of RGC axonal regeneration is also increased by a single application of ciliary neurotrophic factor (CNTF) or fibroblast growth factor 2 (FGF-2) [31]. In the course of that study we found large numbers of cells with the ultrastructural characteristics of macrophages that congregated at the injury site, confirming an observation made by our laboratory two decades earlier [32]. From these results, the question arose as to whether the application of the growth factors could increase the numbers of macrophages present, and it is this question that we address in the present study. The results of these experiments show that application of these growth factors, in particular CNTF, does increase and prolong the numbers of macrophages in the nerve.

## Materials and Methods

### Animals

Adult frogs (*Rana pipiens*) of both sexes were used. They were obtained from commercial sources and kept in tanks with recirculating tap water at 19°C. A total of approximately 50 animals were used for the immunohistochemistry and electron microscopy experiments. All our protocols have been approved by the Institutional Animal Care and Use Committee (IACUC) and follow the recommendations of the Panel on Euthanasia of the American Veterinary Medical Association.

### Surgical Technique for Optic Nerve Crush

With animals under 0.3% tricaine anesthesia, the right eyeball was approached from the palate in which an incision was made; the extraocular muscles were teased aside, and the extracranial portion of the optic nerve was exposed (approx. 3 mm in length). Avoiding large blood vessels, the nerve was crushed at the hallway point using Dumont No. 5 forceps. This leaves the meningeal sheath intact but creates a transparent gap that is completely free of axons. We have confirmed the lack of even the smallest of axons in this region by electron microscopic observation; also, crushing in this manner perturbs RGC survival almost as effectively as cutting [30]. The incision was sutured, and the animals were allowed to recover for several hours in the laboratory under observation before replacing them in their tanks in the animal facility.

### Neurotrophic factor application

Immediately after the optic nerve was crushed, it was placed on a strip of Parafilm and 5 µl of FGF-2 or CNTF solution was applied directly to the crush lesion. The solution was left in place for 5 min, then the Parafilm was removed and the palate sutured. Control applications consisted of 5 µl of phosphate-buffered saline (PBS: 0.1M). For FGF-2 (R & D Systems, MN, USA) and CNTF (Sigma, St Louis, MO, USA) 125 ng total were applied.

### Resin embedding for light and electron microscopy

Animals were euthanized one and two weeks after optic nerve crush and PBS, CNTF, or FGF-2 application (N = 3 - 6 per treatment). With animals under 1% tricaine + 0.04% NaHCO_3_ anesthesia, the head of the animal was cut off and the region of the optic nerve was approached from the palate and left in fixative overnight (2% paraformaldehyde + 2% glutaraldehyde in diluted in 0.1 M cacodylate buffer with 0.05% CaCl_2_). The next day, the samples were washed twice in 0.1 M cacodylate buffer, 5 minutes each. The proximal and distal stumps of the optic nerve and a portion of the eyeball were carefully dissected then postfixed under a fume hood with 1% osmium tetroxide (OsO_4_) diluted in cacodylate buffer for 1 h. Subsequently, the samples were dehydrated in 25%, 50%, and 70% ethanol, 5 minutes each, then were placed in 3% uranyl acetate diluted in 70% ethanol for 1 h. The dehydration process was continued, placing the nerves in 90% ethanol (5 minutes), three times in 100% ethanol (20 minutes each), and 10 minutes in propylene oxide. The nerves were infiltrated with 50/50 Epon-Araldite resin and propylene oxide for 1 h, then in 100% Epon-Araldite and left in the desiccator overnight. The next day the nerve samples were placed in cubic molds and embedded in 100% resin, then placed in a 60°C oven for 24 hours. The resin block was trimmed and, using an ultramicrotome (Sorvall MT-2), transverse sections were cut; semi-thin sections (1 μm thick) for light microscopy, and ultrathin (90 nm) for electron microscopy. For light microscopy, semi-thin sections were stained using methylene blue-azure II and basic fuschin. Thin sections were examined with a JEOL JEM-1011 electron microscope equipped with a Gatan digital camera (Model-832) to describe the ultrastructural features of the nerve.

### Light microscopy cell counts

Serial 1 μm resin sections were cut from the optic nerve, starting at the distal stump and working proximally until the injury site was reached, collecting the sections every 50 μm. The section 100 μm distal to the injury site was used for detailed analysis, because many regenerating axons have reached this point at one week [31]. Composite high magnification light microscope images of the nerve were examined in Adobe Photoshop or GIMP. Color overlays were constructed by a trained observer (JMB), outlining the nerve and macrophage-like cell bodies within it. Image filenames were coded so as to blind the observer to whether they came from experimental or control animals. Macrophage-like cell profiles were identified by their large size, dark cytoplasm, and the presence of granules and/or cytoplasmic vacuoles. The overlays were thresholded in ImageJ (Fiji) and quantified using the Analyze Particles function.

### Immunohistochemistry

After dissection, 3 - 5 eye cups with optic nerves attached were fixed for each control and experimental stage with buffered 2% paraformaldehyde solution for 1 hr. After PBS washing, the tissues were placed in 30% sucrose for cryoprotection at 4°C overnight, and, after being frozen, cryostat sections of 12-20 µm were cut. After air-drying, sections were immersed in 10 mM citrate buffer (pH 6) for 10 min at 60°C. The sections were washed twice (5 min each) in PBS containing 0.3% Triton X-100 + 0.5% bovine serum albumin (BSA) and incubated for 30 minutes in the same buffer containing 10% normal goat serum (NGS; for, ED1) or 10% normal rabbit serum (NRS; for Arg1). They were then incubated with the antibodies against ED1 (1:100; catalog # MCA5709; ABD Serotec) and Arginase 1 (1:100; catalog # sc-18355; Santa Cruz Biotechnology), diluted in 0.1M PBS + 0.3% Triton X-100 + 0.5% BSA, overnight at 4°C. After several washes in the same buffer solution the sections were incubated with goat anti-mouse Cy2 (1:100, Jackson ImmunoResearch Laboratories, Inc.), rabbit anti-goat CY3 (1:100, Jackson ImmunoResearch Laboratories, Inc) for 2 h at room temperature. For ED1 and Arg1 sections, we labeled the cell nuclei with 4’,6-diamidino-2-phenylindole (DAPI) staining for 5 minutes after the secondary antibodies. The sections were rinsed in 0.1 M PBS six times, 5 minutes each, and mounted in Polymount.

Omitting the primary antibodies resulted in the absence of immunostaining. The ED1 antibody recognizes rodent and bovine CD68 or macrosialin, which has some homology with amphibian lysosomal proteins. Additionally, a second polyclonal antibody against CD-68 (Abcam #ab124212) gave a very similar staining pattern. The Arg1 polyclonal antibody recognizes the C terminus of mammalian arginase 1, the likely antigenic regions of which show almost 70% identity with amphibian arginase.

Frozen sections of the whole optic nerve processed with ED1 and Arg1 were used to count the macrophages at the injury site. Alternating longitudinal sections through the nerve were analyzed to avoid overlap of data. The number of stained macrophages of each type was counted from confocal images obtained with a Zeiss Pascal laser scanning confocal microscope, using Zeiss LSM5 Image Browser Software, then expressed as a proportion (Arg1/ED1). The statistical significance was determined using ANOVA with *post-hoc* Tukey-Kramer tests (*P < 0.05, **P < 0.01, ***P < 0.001).

## Results

### Growth factor application increases the numbers of macrophage-like cells in the injured optic nerve

Transverse 1 μm resin sections of control optic nerves were examined with light microscopy (Fig 1). We chose to focus on the region 100 μm distal to the crush site because many regenerating axons have reached this point at one week [31]. Macrophage-like cells were identified by their size, dark staining and granular/vacuolar appearance (Fig 1A and C). In PBS-treated controls, from 71 to105 macrophages-like cell profiles were counted in this region from overlays (Fig 1B, N=6 animals). Because the cross-sectional area of the nerve varied somewhat between preparations (0.089 – 0.225 mm^2^), this count was converted to cell density, giving a mean of 644 ± 38 cells/mm^2^ (N = 6, Fig 1D). Treatment with CNTF increased macrophage-like cell density 2.7-fold to 1751 ± 481 cells/mm^2^ at 1 w (N = 3, Fig 1E). Treatment with FGF-2 was equally effective, increasing cell density 2.5-fold to 1591 ± 309 cells/mm^2^ at 1 w (N = 3, Fig 1E).

**Fig 1.**
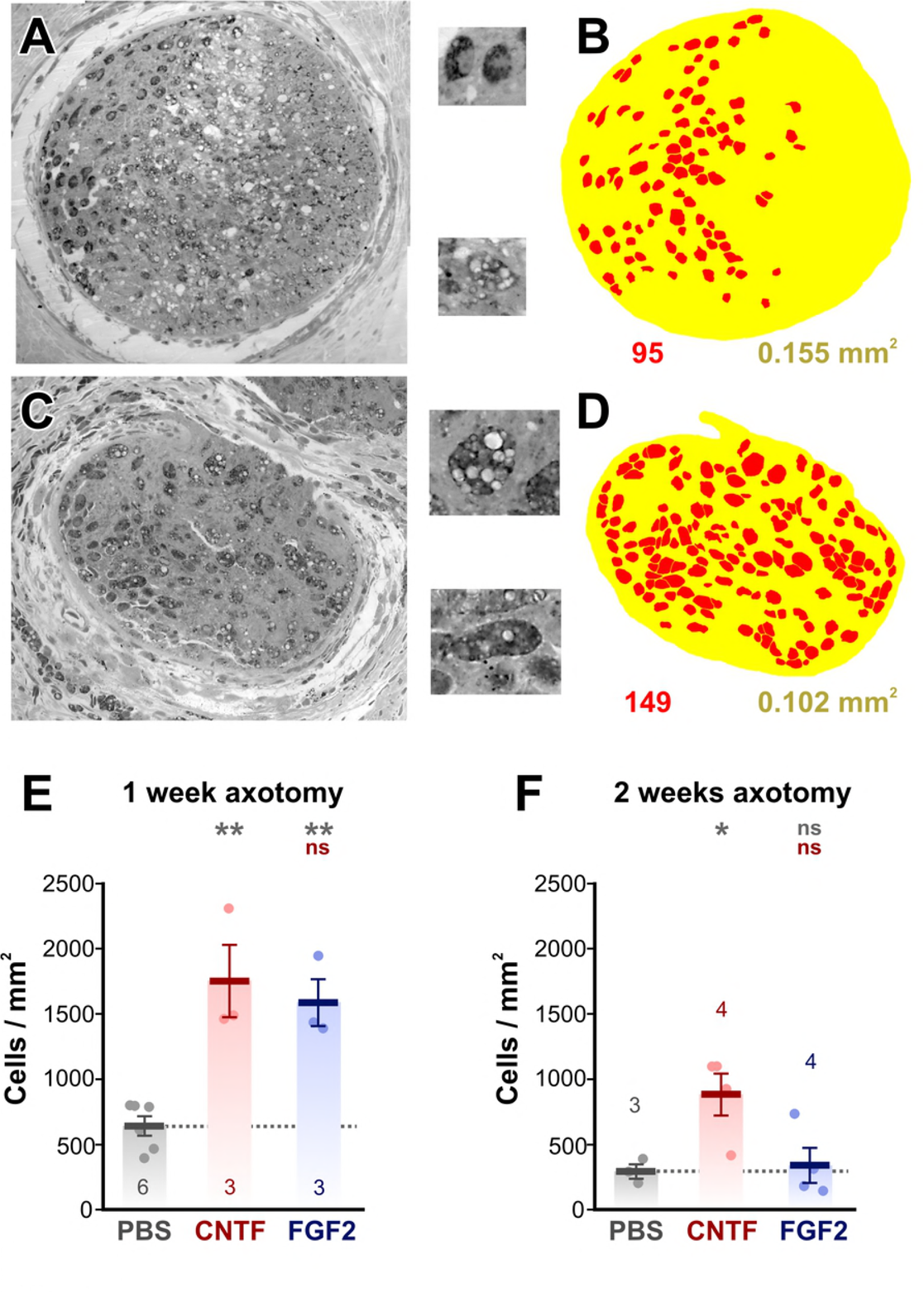
Growth factor treatment increases macrophage numbers. (A and C) Light micrographs of 1 μm resin sections of optic nerves, taken 100 μm distal to the crush zone. A is a PBS-treated control, C is from a CNTF-treated animal. The insets show enlarged examples of macrophage-like cell profiles, showing dark staining, granules, and vacuoles. (B and D) Color overlays of the light micrographs, delineating macrophage-like cell profiles (red) and the nerve itself (yellow). The cell count and nerve area derived from these overlays are shown below. (E and F). Combined scatterplots and barcharts of cell density showing mean ± SEM. Asterisks or “ns” above each column indicate the significance when compared to PBS (row 1, gray) or CNTF (row 2, red) with ANOVA and *post-hoc* Tukey tests. (E) At 1 w after nerve crush there are increases in cell density with CNTF and FGF-2 treatment, compared to PBS-treated controls. (F) 2 weeks after crush, only CNTF treatment shows an effect.

By two weeks after axotomy the numbers of macrophage-like cells in the region 100 μm distal to the crush site was significantly decreased by about half in control PBS-treated animals, to 294 ± 93 cells/mm^2^ (N = 3, p = 0.017, homoscedastic t-test). The cell density remained elevated 3-fold with CNTF treatment at 887 ± 323 cells/mm^2^ (N = 4, Fig 1F), although this was significantly less than at 1 w (p = 0.035, homoscedastic t-test). However, cell density with FGF-2 treatment had fallen to control levels (342 ± 271 cells/mm^2^, N = 4, Fig 1F).

The macrophage overlays allowed the quantification of various parameters of the cell profiles. We were interested to determine whether the cells became larger as a result of growth factor treatment, and so measured their diameter (Feret diameter, ie. longest diameter of each profile). Diameters were segregated in 10 μm bins for each preparation, then these totals were expressed as a percentage of the total number of cells and averaged over the experimental animals (Fig 2). The majority (90%) of the cell profiles fell in the range of 20-40 μm (Fig 2). However, we found no significant changes in cell size as a result of growth factor treatment (ANOVA followed by *post-hoc* Tukey tests), and neither were there any changes in size between 1 week (Fig 2A) and 2 weeks after optic nerve injury (Fig 2B).

**Fig 2.**
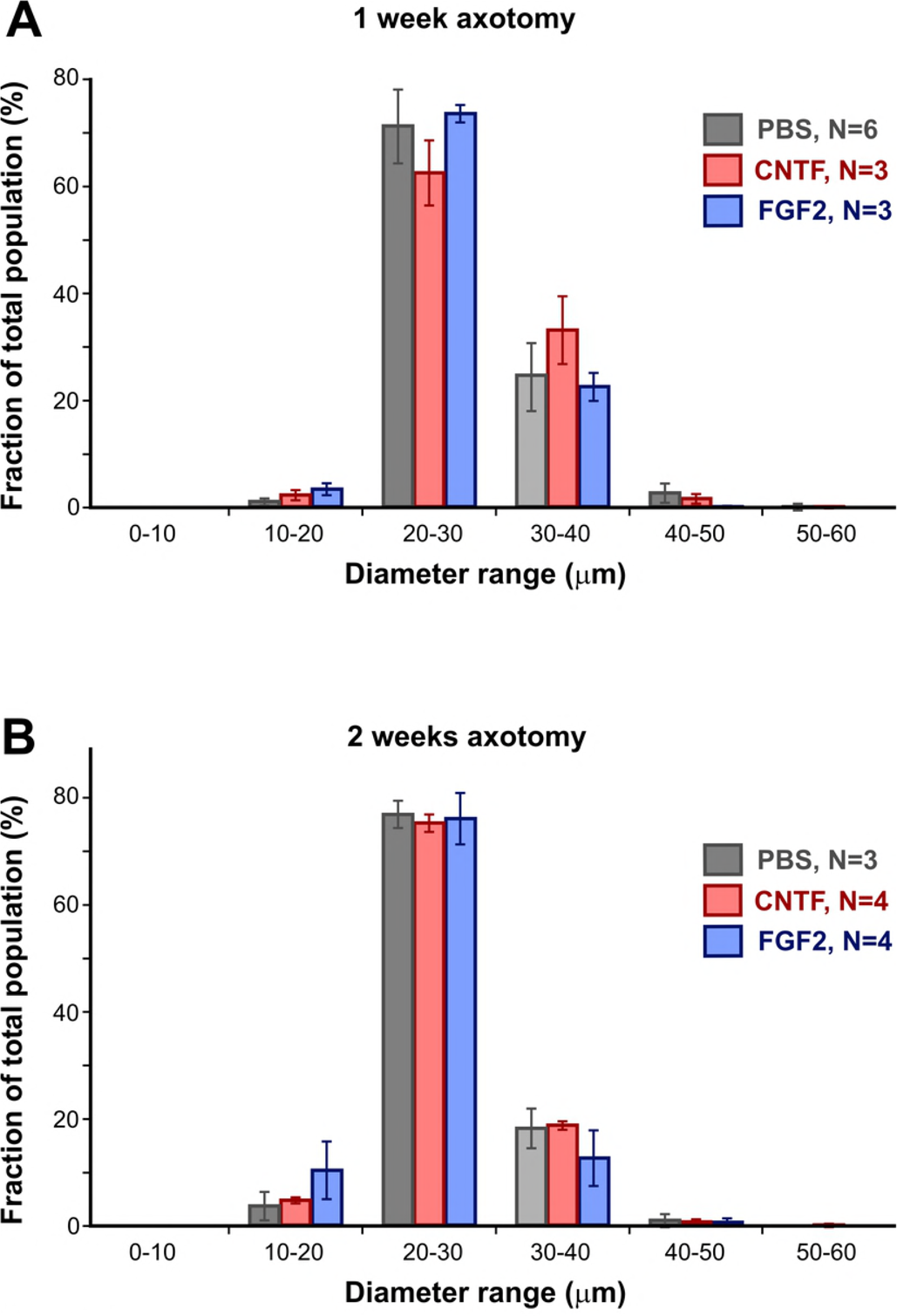
Growth factor treatment does not alter macrophage size. (A and B). Histograms of cell profile diameter (Feret diameter) averaged over several preparations, showing mean ± SEM for each size category. There are no significant differences in cell sizes with growth factor treatment, either at one or two weeks, and no changes in the population of profile diameters between those times.

### Growth factor application does not alter the relative proportions of macrophage types

Macrophages in mammals can be divided into two broad types: pro-inflammatory M1 and alternatively-activated pro-repair M2, which can be distinguished by their expression of different antigenic markers. During spinal cord injury the pro-inflammatory type overwhelms the small, transient, M2 response [16,22]. We were therefore interested to determine whether frog macrophages could be identified as M1 or M2 and to find out whether their relative proportions changed at different times after injury and growth factor treatment.

Longitudinal frozen sections of optic nerves were stained with two antibodies, ED1 and Arg1 (Fig 3). Counterstaining with DAPI showed that most, but not all, cells within the control, PBS-treated optic nerves were ED1-positive (Fig 3A-D). The ED1 antibody recognizes the mammalian CD68 (macrosialin in mouse) protein, which is expressed in the lysosomes of all macrophages [33–35]. BLAST searches indicated some homology with amphibian lysosomal proteins so it is possible that ED1 also labels a *Rana* homolog of CD68. In any case, it is clear from high magnification images (Fig 3A-B) that ED1 does indeed stain lysosome-like structures, confirming its utility as a pan-macrophage marker in amphibia.

**Fig 3.**
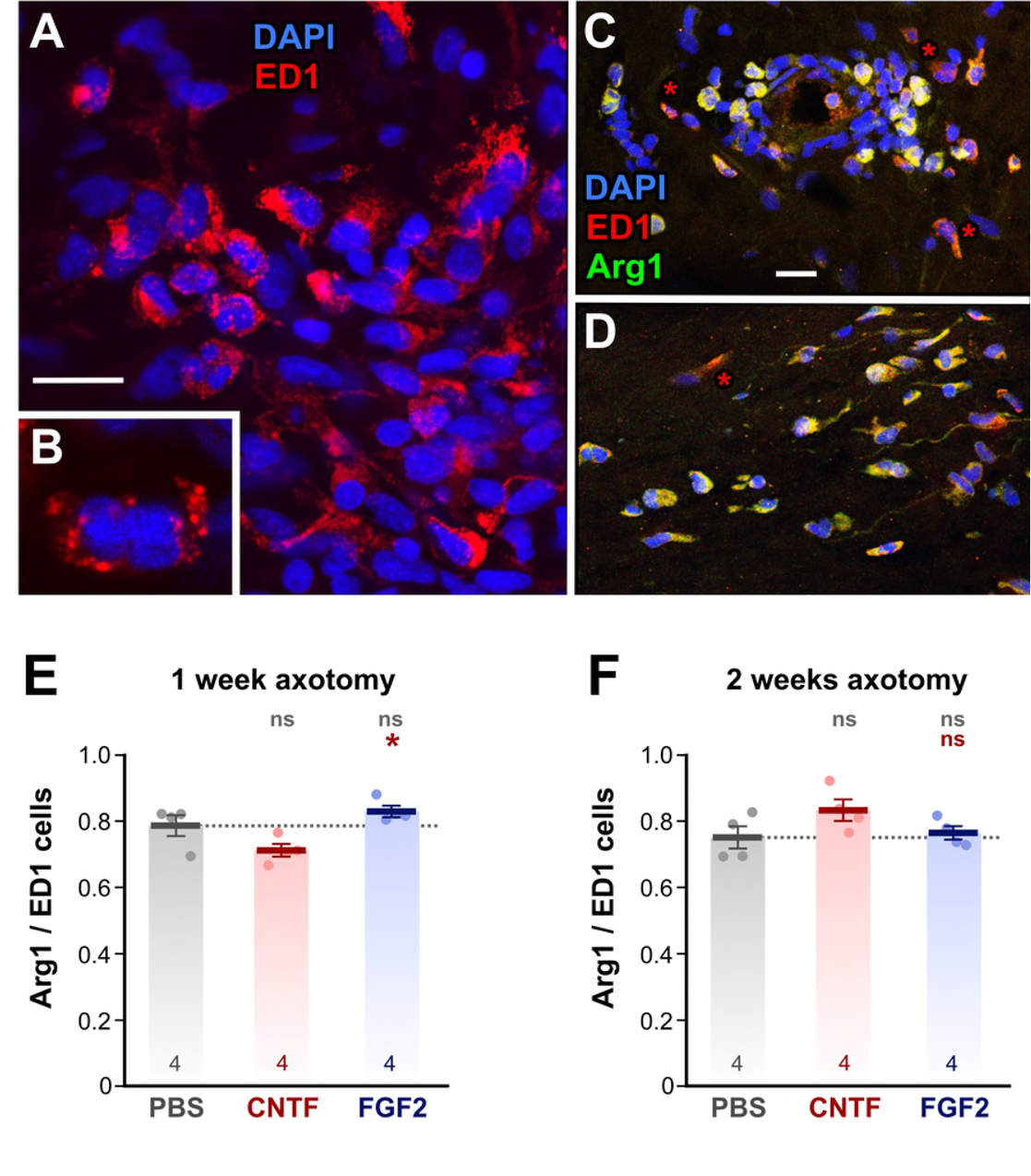
Immunostaining of macrophage subtypes. (A - D) Confocal fluorescence micrographs of immunostained frozen sections of optic nerve 1 week after axotomy. (B) High magnification single slice of a macrophage with an irregular nucleus, clearly showing ED1-stained lysosomes. (C) Double staining of peripherally-located macrophages with ED1 and Arg1. Only a few cells have predominantly ED1 staining with little Arg1 (asterisks). (D) Double staining of centrally-located macrophages with ED1 and Arg1. Only a few cells have predominantly ED1 staining (asterisks). (E and F). Combined scatterplots and barcharts of Arg1/ED1 ratio showing mean ± SEM. Asterisks or “ns” above each column indicate the significance when compared to PBS (row 1, gray) or CNTF (row 2, red) with ANOVA and *post-hoc* Tukey tests. At 1 w (E) and 2 w (F) after nerve crush there is no change in the Arg1/ED1 proportion with CNTF and FGF-2 treatment, compared to PBS-treated controls. Scale bar: 20 μm in A, C, D; 10 μm in B.

The Arg1 antibody recognizes the C terminus of mammalian arginase 1, which is a classical marker for pro-repair M2 macrophages [36]. A BLAST search of the likely antigenic regions of this molecule showed an almost 70% identity with amphibian arginase, making it likely that Arg1 stains arginase, and therefore M2-like macrophages, in *Rana*. We carried out double immunostaining of optic nerve sections using Arg1 along with ED1 to determine how the relative proportion of putative M2 macrophages changed after injury and growth factor treatment (Fig 3). In fact, we found that at 1 week after injury the proportion of Arg1/ED1 macrophages was approximately 80% (Fig 3E), and that it remained at this level at 2 weeks after injury (Fig 3F). Treatment with CNTF or FGF-2 had no effect on the relative proportions of Arg1/ED1 macrophages, even though the total numbers were increased (see Fig 1). This result indicates that in *Rana*, unlike rodents, there is a high proportion of putative M2 (pro-repair) macrophages present soon after injury, and that this proportion remains unchanged for at least 2 weeks, during the period when regeneration is taking place [31].

### Ultrastructural characteristics of macrophages in the optic nerve after injury

The optic nerve is surrounded by the meningeal sheath, a connective tissue layer that protects the CNS and separates it from the environment. Beneath this is the *glia limitans*, a layer made up of astrocytes, glial cells that characteristically exhibit large bundles of intermediate filaments and desmosomes [32]. One week after optic nerve crush, actively phagocytosing macrophages were present inside the optic nerve, presumably playing a role in clearance of the debris (Fig 4). Cells with large vacuoles were common, particularly in central regions of the nerve (Fig 4D). These vacuoles appeared to contain the remnants of degenerating axons and myelin. Two weeks after nerve crush, there appeared to be fewer macrophages in the nerve (Fig 4F, G). Those which were present, both peripherally and centrally, contained fewer vacuoles with axonal debris and more pale inclusions which were likely composed of lipids, since they had no bounding membrane (Fig 4F inset).

**Fig 4.**
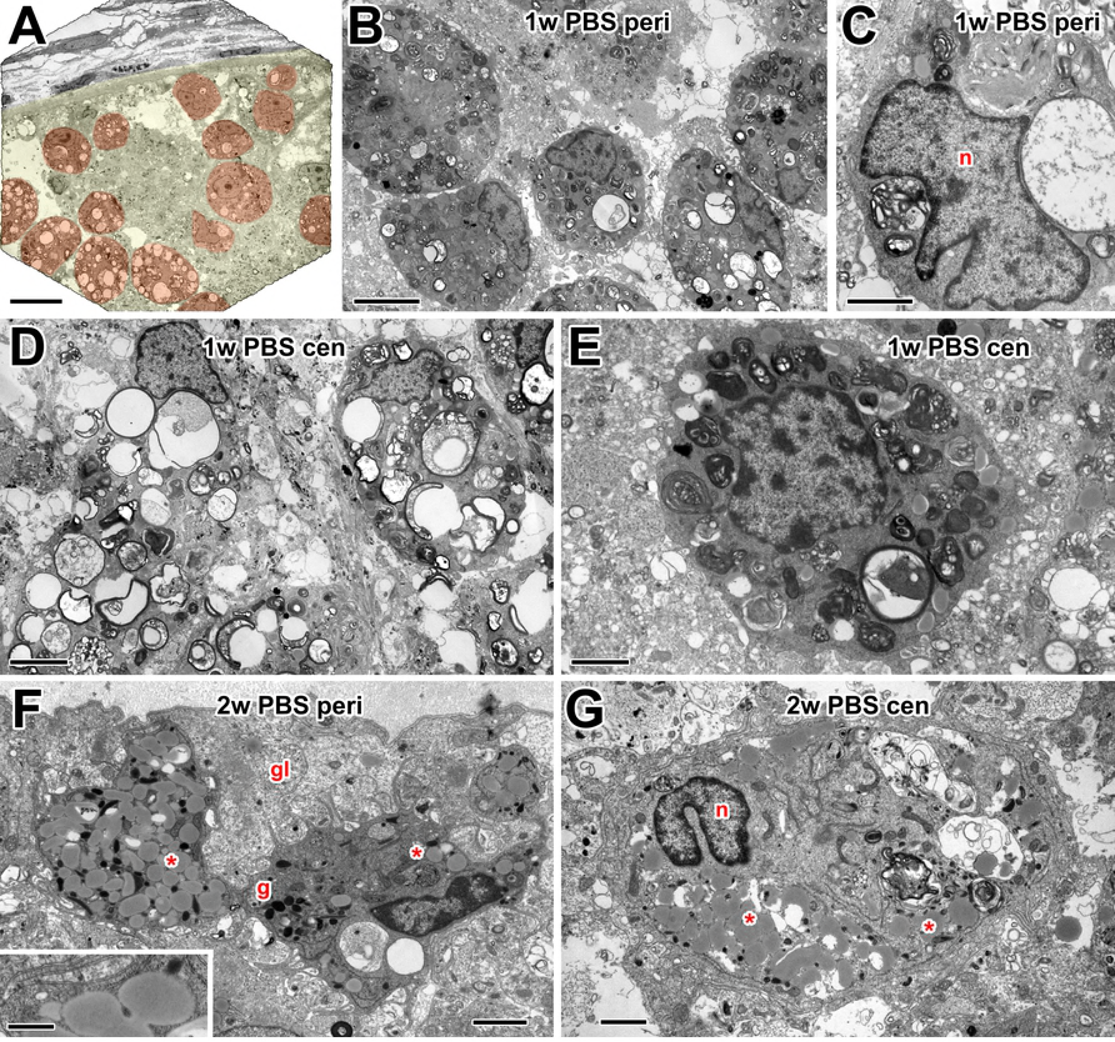
Electron microscopy of macrophages in PBS controls. (A) Low power electron micrograph of the optic nerve periphery at 1 w after injury, with a pale yellow overlay indicating the nerve and pale red overlay indicating numerous macrophages within it. (B) Peripheral macrophages containing a variety of residual bodies, products of phagocytic activity. (C) Macrophage with an irregular nucleus (n) and large debris-filled residual bodies. (D) Centrally located macrophages containing large vacuoles with apparent remnants of degenerating axons. (E) Central, smaller, macrophage/microglia with denser residual bodies. (F) Peripheral macrophages at 2 weeks, located within the glia limitans (gl). Some vacuoles with axonal debris are present but the cytoplasm also contains lipid inclusions (asterisks) and dark granules (g). Inset: high magnification view of putative lipid inclusions, showing lack of a bounding membrane. (G) Large central macrophage with a bilobed nucleus (n) at 2 weeks, containing a mixture of vacuoles with axonal debris, and lipid inclusions (asterisks). Scale bar: 20 μm in A; 5 μm in B, D; 2 μm in C, E, F, G; 0.5 μm in inset.

### CNTF treatment increases optic nerve macrophage activity

One week after optic nerve injury and treatment with CNTF, large numbers of macrophages were found within the optic nerve, both peripherally (Fig 5A, B) and centrally (Fig 5C, D). These appeared to be highly active phagocytically, judging by the large numbers and sizes of debris-containing vacuoles and residual bodies. Additionally, other cell types, such as neutrophils (Fig 5C) are also present. Compared to PBS controls, there was the appearance of more macrophages with large vacuoles at the periphery of the nerve (Fig 5A, B).

**Fig 5.**
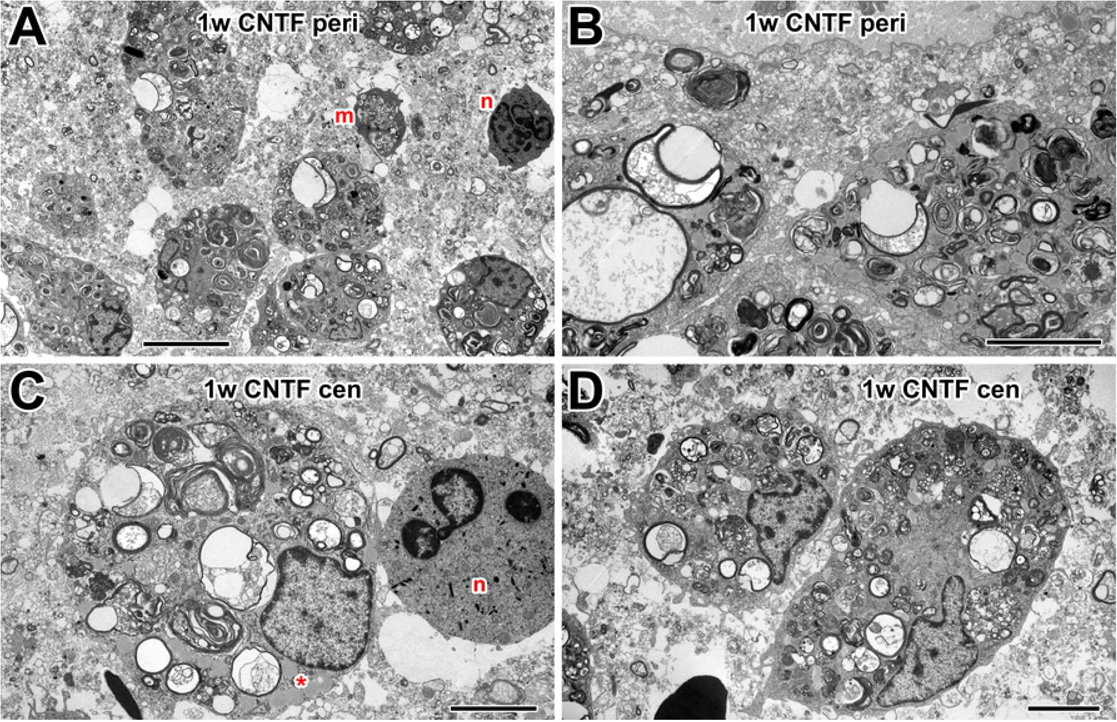
Electron microscopy of macrophages in 1 w CNTF-treated nerve. (A) Low power electron micrograph of the optic nerve periphery, showing numerous macrophages containing large debris-filled vacuoles. Also present are smaller microglia (m) and neurotrophil-like cells (n). (B) Active macrophages, with many large vacuoles, at nerve periphery. (C) Small, central macrophage/microglia with multiple lamellar residual bodies and also lipid inclusions (asterisk). Next to it is a neutrophil-like cell (n). (D) Centrally located macrophages with numerous residual bodies. Scale bar: 10 μm in A; 5 μm in B, C, D.

Two weeks after optic nerve injury and CNTF treatment, large numbers of cells were still present within the optic nerve, both peripherally (Fig 6A-C) and centrally (Fig 6D-F). Some were very large (Fig 6D) and probably were indeed macrophages, while others were smaller in size (Fig 6E) and may represent microglia. Compared to PBS controls at this stage, the more numerous cells in CNTF-treated nerves still showed signs of ongoing phagocytosis, containing vacuoles with axonal debris and multilamellar myelin/membrane remnants, as well as the numerous lipid inclusions seen at 2 w in PBS controls. Some of these cells also contained small dark structures that resemble secretory granules (Fig 6A, D).

**Fig 6.**
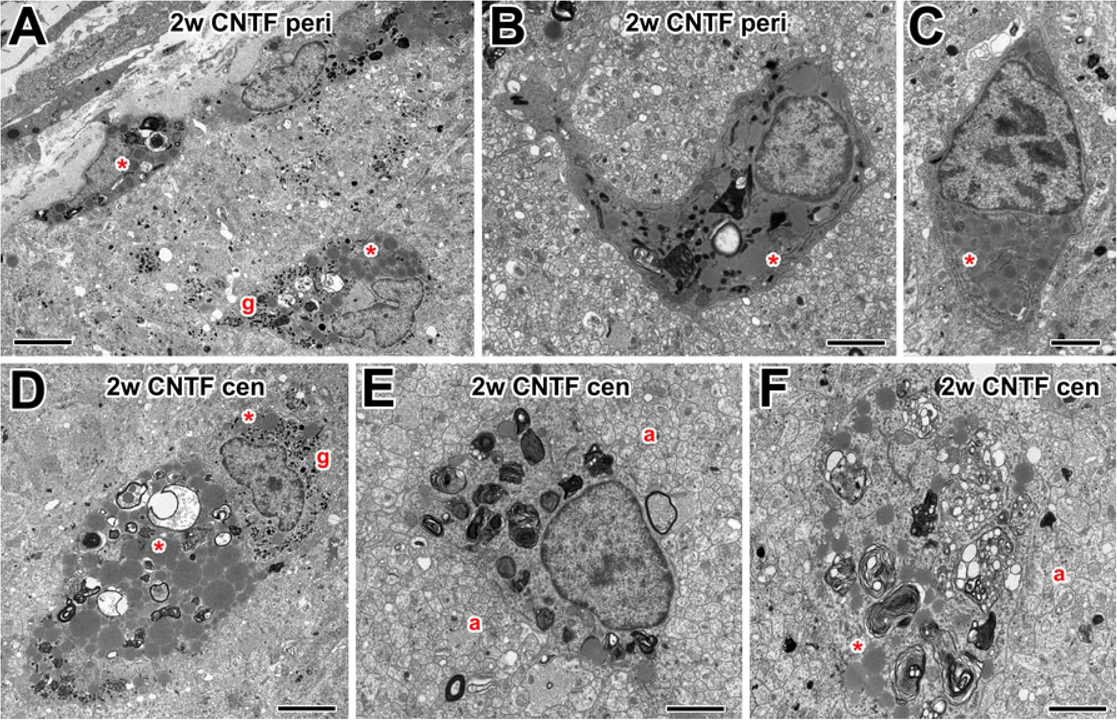
Electron microscopy of macrophages in 2 w CNTF-treated nerve. (A) Low power electron micrograph of the optic nerve periphery, showing macrophages containing some residual bodies, also lipid inclusions (asterisks) and small dark granules (g). (B) Peripheral microglia-like cell containing residual bodies and lipid inclusions (asterisk). (C) Peripheral unknown cell type containing rough endoplasmic reticulum and lipid inclusions (asterisk). (D) Large, centrally located macrophage with residual bodies, vacuoles containing axonal debris, and lipid inclusions (asterisks). Around its nucleus are numerous dark granules (g). (E) Central macrophage/microglia with myelin debris, in close proximity to many small axons (a). (F) Macrophage with residual bodies and lipid inclusions, in proximity to axons (a). Scale bar: 5 μm in A, D; 2 μm in B, C, E, F.

### Effects of FGF-2 treatment on macrophage ultrastructure

One week after optic nerve injury and treatment with FGF-2, as with CNTF, large numbers of active macrophages were observed in the optic nerve (Fig 7A, B). These contained vacuoles with axonal debris and lamellar residual bodies. Other types of blood cells, such as eosinophils, were occasionally found (Fig 7A). At two weeks after FGF-2 treatment, fewer macrophages were present in the optic nerve, similar to controls. Those that were found were large, contained numerous lipid inclusions (see above) (Fig 7C-E), and some had residual bodies and dark (possibly secretory) granules (Fig 7D).

**Fig 7.**
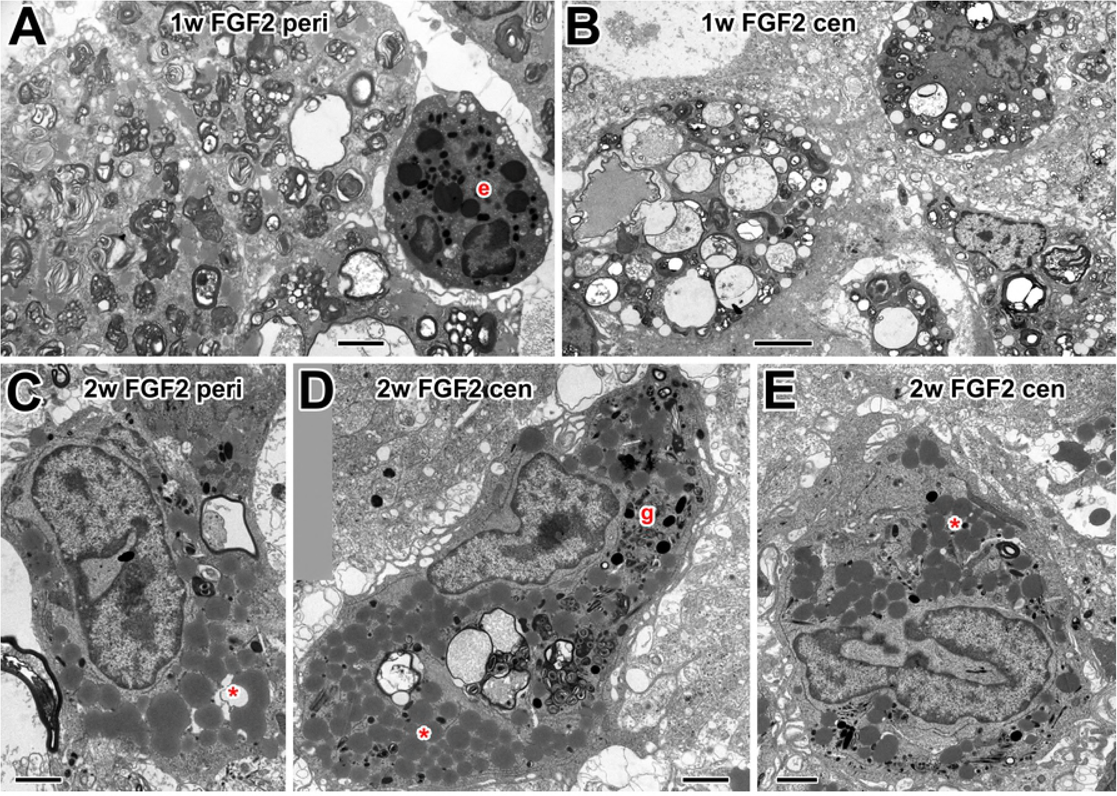
Electron microscopy of macrophages in FGF-2-treated nerve. (A) Peripheral macrophages at 1w, containing lamellar residual bodies. Also present is a possible granulocyte, with a bilobed nucleus and dark granules (e). (B) Macrophages in the central optic nerve at 1w, containing vacuoles with axonal debris. (C) Peripheral nerve at two weeks showing macrophage with lipid inclusions (asterisk). (D) Large, centrally located macrophage at 2 w with residual bodies, axonal debris and lipid inclusions (asterisks). Around its nucleus are numerous dark granules (g). (E) A similar large central macrophage with a highly indented nucleus and lipid inclusions (asterisk). Scale bar: 2 μm.

## Discussion

Early work from this laboratory and others noted the appearance of macrophages in the axotomized optic nerve of the frog *Rana pipiens* [32,37]. However, this present study is the first detailed characterization of those macrophages, investigating the timing of their appearance after axotomy and their heterogeneity of their subpopulations. In addition, here we also investigate how the growth factors CNTF and FGF-2, which we have shown to affect the speed and the number of regenerating axons [31], also affect the macrophage populations during optic nerve regeneration.

Light microscope counts of macrophage-like cell profiles showed the appearance of many cells in the regenerating region by 1 week after injury, the numbers of which subsequently declined by half at 2 weeks. This result is consistent with our earlier qualitative report [32], in which the nerve was cut and the stumps separated. Macrophages in other animals show similar large yet transient influxes after nerve injury, for example rat sciatic nerve [38], axolotl spinal cord and peripheral axons [39], and the optic nerves of rat, goldfish and *Xenopus* tadpoles [40–42].

We found that application of CNTF to the lesion doubled the number of macrophages in the nerve without affecting their size distribution, and that this effect persisted for 2 weeks after injury, long after the CNTF itself would had disappeared. There are few studies which have investigated the possibility of such a chemoattractive effect *in vivo*, although in the rat eye CNTF injection does increase the recruitment of blood-derived macrophages [43]. In addition, CNTF elicits concentration-dependent chemotaxis of macrophages *in vitro* [44]. We also found that FGF-2 application increased the number of macrophages in the nerve, but the effect was temporary, and had disappeared by 2 weeks after injury. There are no comparable studies of FGF-2 on macrophages in other systems, however it has been shown that it increases the migration and survival of tumor-associated macrophages [45]. On the contrary, however, there appears to be increased macrophage activity after sciatic nerve injury in FGF-2-knockout mice [46].

In our previous study we investigated the effect of a single application of growth factor to the injury site on regenerating axons at 2 weeks after injury [31]. We showed that axon speed was increased 138% by CNTF and 63% by FGF-2, while axon numbers were increased 72% by FGF-2 and 52% by CNTF [31]. These prolonged effects must have far outlasted the immediate actions of the factors themselves and presumably resulted from longer-term changes in the axonal microenvironment. In the present study we investigate one such change, i.e. the increase in macrophages, and show that both CNTF and FGF-2 greatly increase the numbers of these cells, but only at 1 week after injury. The timing of these events suggests a causative effect - increased numbers of macrophages results in an enhancement of axonal growth. However, we need to test this hypothesis by inhibition of macrophages, and this will be the focus of a future study.

By what means do the macrophages enhance axonal regrowth? One obvious answer from our electron microscope observations is that they phagocytose the distal stumps of severed axons, thus removing a physical impediment to axonal elongation, which otherwise could remain in place for up to 3 months after axotomy [47]. Another possibility is that phagocytosis removes an inhibitory chemical barrier in the myelin, as in the mammalian CNS [48]. However, *Xenopus* optic tract myelin is not inhibitory to axonal extension (unlike that from the spinal cord)[49], despite both expressing an ortholog of Nogo [50]. It is more likely that, as in fish [3,51,52], the relatively few frog oligodendrocytes and their myelin are not inhibitory.

At 1 week after injury, irrespective of growth factor treatment, most macrophages were observed to have large vacuoles containing what appeared to be axon fragments, and multilamellar bodies that may represent myelin debris. By 2 weeks, in PBS- and FGF-2-treated preparations the remaining macrophages tended to contain what appeared to be lipid inclusions (not bounded by a membrane). Similar lipid inclusions, which probably represent the end points of myelin and membrane destruction, are seen in *Xenopus* astrocytes during the extensive myelin remodeling that takes place during metamorphosis [53]. More significantly, they are also the hallmark of the so-called “foamy” macrophages seen in the injured mouse spinal cord after the accumulation of excessive myelin debris [54]. These PBS- and FGF-2-treated macrophages may therefore be in the final stages of phagocytosis, while in CNTF-treated nerves the more numerous macrophages still showed signs of ongoing phagocytosis, in addition to the lipid inclusions. This apparent extension of the active phagocytic period, along with the overall higher numbers of macrophages, appears to be beneficial to axonal extension. Another possibility is that the cells themselves secrete growth-promoting substances, as is the case with mouse M2 macrophages [22], which have increased expression of, for example, insulin-like growth factor 1 and 2 and hepatocyte growth factor [55].

Early studies gave rise to ambiguous conclusions regarding the beneficial or harmful effects of macrophages on CNS repair after injury. However, in 2009 it was demonstrated that the entry of different macrophage subsets into the injured rat spinal cord, either “classically activated” proinflammatory (M1) or “alternatively activated” anti-inflammatory (M2) [56], had different effects, the former being neurotoxic while the latter promoted regeneration [22]. The predominance of the M1 type over the M2 was postulated to be one of the factors responsible for poor CNS regenerative capabilities [16,22]. Macrophage polarization into M1/M2-like phenotypes, although undoubtedly a somewhat oversimplified classification [57], appears to be conserved throughout the vertebrates [58]. One of the distinguishing traits of the M2-like phenotype is the expression of arginase, which is purported to have healing functions [59]. Using double immunostaining of a putative pan-macrophage marker (ED1) along with an antibody against arginase (Arg1), we found that the majority (80%) of macrophages that entered the crushed frog optic nerve were Arg1+, and therefore presumably of the M2 phenotype.

Application of CNTF or FGF-2 greatly increased the overall macrophage numbers without altering the proportion of M2-like cells. Since we have shown previously that both CNTF and FGF-2 increase the numbers and speed of axons that are regenerating through the frog nerve [31] it is clear that the increased presence of M2-type macrophages is entirely consistent with them having a beneficial effect on axonal regrowth, similar to that proposed in rat [22].

Phagocytosis is, by definition, the morphological hallmark of the macrophage; however, it is not clear from the literature whether macrophage polarization is related to phagocytic activity. M1 macrophages in wounds have been described as highly phagocytic, in contrast to the M2 phenotype which accelerate wound closure [60]. On the other hand, in recent *in vitro* assays, it was shown that macrophages activated by IL-4 or IL-10 (equivalent to M2) showed higher levels of phagocytic activity compared to those activated by IFN-⇘ (equivalent to M1) [61,62]. Our electron microscope results clearly show intense phagocytic activity in all of the macrophages that we examined, particularly at 1 week after axotomy, with or without growth factor treatment. Since we estimate that 80% of frog macrophages are Arg1-positive, and thus correspond to the M2 subtype, our results support the idea that M2 macrophages *in vivo* are phagocytically active, and that this activity is beneficial for axonal regrowth.

Our current work shows that the macrophage population in the frog optic nerve is more diverse than previously thought. This system, in which the optic nerve can regenerate, provides a good model to further study the roles of these populations in modulating axonal regeneration and ganglion cell survival.

## Supporting Information Captions

**S1 Table. Light microscope cell analysis data.**

**S2 Table. Immunostaining analysis data.**

